# Functional clusters for shape, texture, and motion encoding in macaque V2

**DOI:** 10.1101/2025.11.01.686029

**Authors:** Taekjun Kim, Rohit Kamath, Gaku Hatanaka, Tomoyuki Namima, Celeste Dylla, Wyeth Bair, Anitha Pasupathy

**Affiliations:** Department of Neurobiology & Biophysics and Washington National Primate Research Center, University of Washington, Seattle, WA 98195; Graduate School of Frontier Biosciences, The University of Osaka and Center for Information and Neural Networks, National Institute of Information and Communications Technology, Suita 565-0871, Japan

## Abstract

Macaque primary visual cortex (V1) exhibits exquisite columnar organization, while midlevel area V4 does not. Here we investigated the functional organization and representational bases of intervening area V2 with high-density Neuropixels recordings and a variety of visual stimuli—shape, texture, drifting grating, and translational motion patches. We observed dense clusters of similarly tuned neurons often spanning ∼500 µm for shape and motion stimuli, and larger for texture stimuli, consistent with a columnar structure. In terms of representational bases, V2 responses were largely explained by stimulus features based on local image statistics: shape tuning is well-modeled by a linear combination of orientation filters, and direction selectivity is stronger with surface compared to object motion, in striking contrast to V4. Overall, our results support the progression from columns to sparse clusters as neuronal representations transform from encoding local features and feature conjunctions in V1/V2 to a high-dimensional object-based code in V4.

**Significance Statement:** By recording hundreds of neurons simultaneously across layers of macaque visual area V2, we show the first evidence of exquisite fine-scale functional clusters that encode higher-order shape, texture, and motion features, extending well beyond the classic orientation-selective columns seen in area V1. Comparative analyses with V4 further reveal distinct representational bases and organization patterns between adjacent cortical areas, offering insights into how columnar organization is preserved in early visual areas (V1 and V2) but markedly attenuated in higher-order cortex such as V4.

## Introduction

The second stage of cortical visual processing in the primate, area V2, occupies roughly 10% of macaque neocortex (Vanni et al., 2020) and is critical for processing both visual form and motion information. Studies have demonstrated selectivity for local orientation and spatial frequency (Foster et al., 1985; Gegenfurtner et al., 1996; Felleman et al., 2015), as in primary visual cortex (V1), but also for higher-order texture statistics (Freeman et al., 2013; Ziemba et al., 2016; Okazawa et al., 2017), stimulus form (Hegdé and Essen, 2000; Ito and Komatsu, 2004; Anzai et al., 2007), border ownership (Zhou et al., 2000) and stimulus motion contrast (Hu et al., 2018; Ma et al., 2021). Past studies using optical imaging methods have demonstrated remarkable organization across the cortical surface for features such as orientation, color, and motion (Ts’o et al., 1990; Lu and Roe, 2008; Felleman et al., 2015). Here we used high-density probes to study sub-populations of V2 neurons across laminae with a broad range of stimulus features of texture, form, and motion. Our studies are motivated by two goals. First, we wish to gain deeper insights into the representational bases and functional organization of V2. Prior work has demonstrated a dramatic shift in terms of functional organization and representational bases from macaque V1 to V4: columnar organization in V1 (Hubel and Wiesel, 1974; Obermayer and Blasdel, 1993) to sparse clusters in V4 (Namima et al., 2025), and basic feature detection in V1 (De Valois et al., 1982) to object-based encoding in V4 (Pasupathy et al., 2020).

Revealing the representational bases and organization in the intervening area V2 for shape, texture, and motion stimuli will advance our understanding and constrain models of the ventral stream. Second, we aim to contribute to the debate on the origin and significance of columns (Horton and Adams, 2005). Some studies have posited that columns represent a critical functional unit (Mountcastle, 1957; Hawkins et al., 2017); on the other hand, demonstrations of their absence in some species (Girman et al., 1999; Adams and Horton, 2006) and in some brain regions (Johnson et al., 2000; Sato et al., 2009) raise the possibility that columns may instead arise as a by-product of representational expansion (Ringach, 2007; Jang et al., 2020), i.e., a small number of inputs project onto a large number of neurons. Moreover, columnar structure may also be less likely to develop when converging afferents exhibit high dimensionality (Purves et al., 1992). Our recent observation in area V4 (Namima et al., 2025), where shape and texture selectivity appeared less spatially clustered, supports this latter hypothesis and prompts further investigation into whether similar patterns exist in V2.

We used high-density multi-contact probes (Neuropixels, IMEC) to simultaneously record responses from dozens of neurons in area V2 of anesthetized macaque monkeys. We used a broad range of stimuli to study responses—2D shape silhouettes, naturalistic textures and their spectrally-matched noise counterparts, sinusoidal drifting gratings and second order translational motion stimuli over a range of spatial displacements. To identify functional clusters of similarly tuned neurons, we first measured the similarity in tuning for each stimulus class by calculating the response correlations between all pairs of simultaneously recorded neurons and then performed cluster analyses. Comparison with previously collected data from area V4 using identical stimuli (Bigelow et al., 2023; Namima et al., 2025) reveals a striking distinction between the two areas in terms of functional organization and representational basis. V2, but not V4, exhibits extensive clustering for neurons encoding similar local features of an image. We also observed differences between V2 and V4 in terms of the representational basis for shape and motion stimuli. Additionally, our large-scale dataset offers new insights into the temporal and laminar dynamics of texture selectivity, highlighting distinct patterns in the processing of perceptual texture dimension of coarseness, directionality, and regularity.

## Materials and methods

### Animal preparation

Electrophysiological recordings were conducted in visual area V2 of three anesthetized, paralyzed rhesus macaques (*Macaca nemestrina*: a 3-year-old 6.7 kg male, a 9-year-old 6.6 kg female, and a 5-year-old 5.5 kg female). The positioning of the craniotomy was guided by stereotaxic coordinates based on standard macaque brain atlases and confirmed by identifying cortical vasculature and sulcal landmarks after durotomy. This placement allowed access to both primary visual cortex (V1) and area V2. To accurately identify the border between areas V1 and V2, we characterized the ocular dominance structure using the widefield optical imaging technique described below. During widefield imaging, the exposed cortex was covered with a glass coverslip to stabilize the surface and minimize optical distortion. For subsequent electrophysiological recordings, the cortex was instead covered with an artificial dura to protect the tissue.

Experiments typically lasted four to five days. Anesthesia was maintained throughout the experiment using a continuous infusion of sufentanil citrate (4-6 μg/kg/hr) and neuromuscular blockade was induced with vecuronium bromide (0.1 mg/kg/hr). Both medications were administered intravenously in lactated Ringer’s solution (8 ml/kg/hr) supplemented with 2.5% dextrose. Throughout the experiments, artificial respiration was maintained, with the respiratory rate adjusted to keep expired carbon dioxide levels within a physiological range (3.8-4.0%). Body temperature was maintained at approximately 37°C using a heating pad.

### Intrinsic optical signal imaging

In two monkeys, we conducted intrinsic optical signal imaging to identify the V1/V2 border with the ocular dominance pattern in V1. A square-wave drifting grating (4 s, 1.0 cycles/deg, 4 Hz) was presented on the display (Display++, Cambridge Research Systems) at a distance of 127 cm. Eight directions were randomly presented in each session for 20 repeats for each condition. One of the eyes was occluded with a shield plate. The image was acquired with sCMOS camera (Zyla-4.2 USB3, Andor) equipped with a tandem lens configuration (two Nikkor 50 mm, Nikon) to achieve 13.3 × 13.3 mm Field of View. A glass coverslip was placed on the cortex to prevent motion artifacts. The cortex was illuminated with an LED at 625 nm wavelength (M625L4, Thorlabs). Image acquisition was controlled by Micro-manager software (μManager, https://micro-manager.org) (Edelstein et al., 2014). The response amplitude was calculated as a fraction of the image intensity from the baseline. First, baseline intensity R0 was calculated as an average over one second before the stimulus onset. Then the driven response (R1) was calculated as an average between five and six seconds after the stimulus onset. Finaly, the response amplitude was calculated as difference over baseline (R1-R0)/R0. The ocular dominance pattern was visualized as a *t*-statistic between the left eye condition and the right eye condition. The vasculature pattern was also acquired with the same imaging setup under 530 nm illumination (M530L4, Thorlabs) to guide electrode penetration location.

### Electrophysiology

Extracellular recordings were carried out using Neuropixels 1.0 and Neuropixels 1.0 NHP (IMEC). Neural signals from the probe and non-neural event signals from other devices (sync pulse generator, photodiode) were acquired using a PXIe acquisition module (PXIe_1000, IMEC) and multifunction I/O module (PXI-6224, National Instruments-NI).

These were transmitted to a data acquisition Windows computer via the PXI Chassis (PXIe-1071, NI). Action potential (AP) and local field (LF) potential signals from the 384 probe contacts (AP: 30 kHz sampling rate; LF: 2.5 kHz sampling rate) were amplified and bandpass filtered (AP: 0.3-10 kHz; LF: 0.5-500 Hz), and stored for offline analysis (spikeGLX, https://billkarsh.github.io/SpikeGLX/). Square-wave sync pulses generated with a microcontroller board (Arduino R3, Arduino) were stored in both the neural and non-neural event data recording streams so that spike timing and photodiode detection could be synchronized in offline analysis.

For probe insertion, we made a small slit (2–3 mm wide) in the artificial dura and passed the probe through it. The exposed cortical surface was maintained by the application of artificial cerebrospinal fluid to prevent drying. Probes were inserted using an oil hydraulic micromanipulator (Narishige, MO-97A) until cortical penetration was achieved and neural activity was detected. Additionally, a custom-built presser foot was used to provide tension to the cortical surface, providing additional stability during probe insertion.

For each individual recording site, binary data files collected across experiments were combined into a single binary file. Combined binary files were then processed with open-source automated spike sorting software (Kilosort 2.5/3) (Pachitariu et al., 2024) with default parameters. The output was subsequently manually curated using the Phy2 interface (https://github.com/cortex-lab/phy) to ensure the quality of single-unit isolation. The detailed procedures for this curation are described elsewhere (Bigelow et al., 2023; Namima et al., 2025). Only well-isolated single units were included in the data analysis.

Following automated and manual spike sorting, a final preprocessing step was performed to align the different data recording streams. Because neural and non-neural event signals were recorded on separate streams with independent clocks, sync pulse edge times were linearly regressed to bring spike times, event timestamps, and photodiode signals into a common temporal reference frame. Subsequent analyses were then performed using custom Python scripts.

### RF characterization

Each penetration began with a hand mapping procedure to roughly identify the spatial RFs of neurons recorded along the length of the probe. This preliminary RF mapping was performed for 2–4 channels that were well spaced along the probe’s length.

To precisely characterize the RF locations, we conducted an automated mapping procedure on a 7 × 7 (or 9 × 9) grid (1° step) centered on the RF estimated by hand-mapping. For this experiment, we used a random dot motion patch (diameter, 1.4°) and presented the stimulus for 300 ms, 10 times at each of the 49 (or 81) grid locations in random order. The RF center was then estimated by fitting a 2D Gaussian to the measured responses (for more details, see (Kim and Pasupathy, 2024)). Based on prior studies, we estimated the V2 receptive field size as (RF diameter = 0.5 + 0.3125 × RF eccentricity) (Gattass et al., 1981). During each recording session, visual stimuli were centered on the average of the estimated RF centers across all channels.

### Visual stimuli

During each session, we presented shape, texture, drifting grating and translational motion patch stimuli in separate blocks. Each stimulus was presented 10 times in random order.

All stimuli were presented against an achromatic background (17 cd/m^2^. x=0.33 and y=0.33 in CIE-xy chromaticity coordinates).

#### Texture stimulus set

The texture stimulus set was composed of 40 naturalistic grayscale textures, drawn from eight groups (five exemplars per group) defined by three perceptual dimensions (i.e., coarseness, directionality, regularity) known to affect human texture perception (Figure 1A). The quantification methods have been described in detail previously (Kim et al., 2019, 2022); briefly, coarseness and directionality indices, computed from the Fourier power spectrum of the images, capture spatial frequency distribution and orientation bias of each texture, respectively. Regularity was computed from the 2D auto-correlation map and reflects the repetitiveness of the texture pattern. Textures were grouped based on high and low values along each of these attributes (see diagrams in Figures 1 & 6), and neuronal responses were compared between the groups (e.g., coarse vs. fine, directional vs. non-directional, regular vs. Irregular).

**Figure 1.**
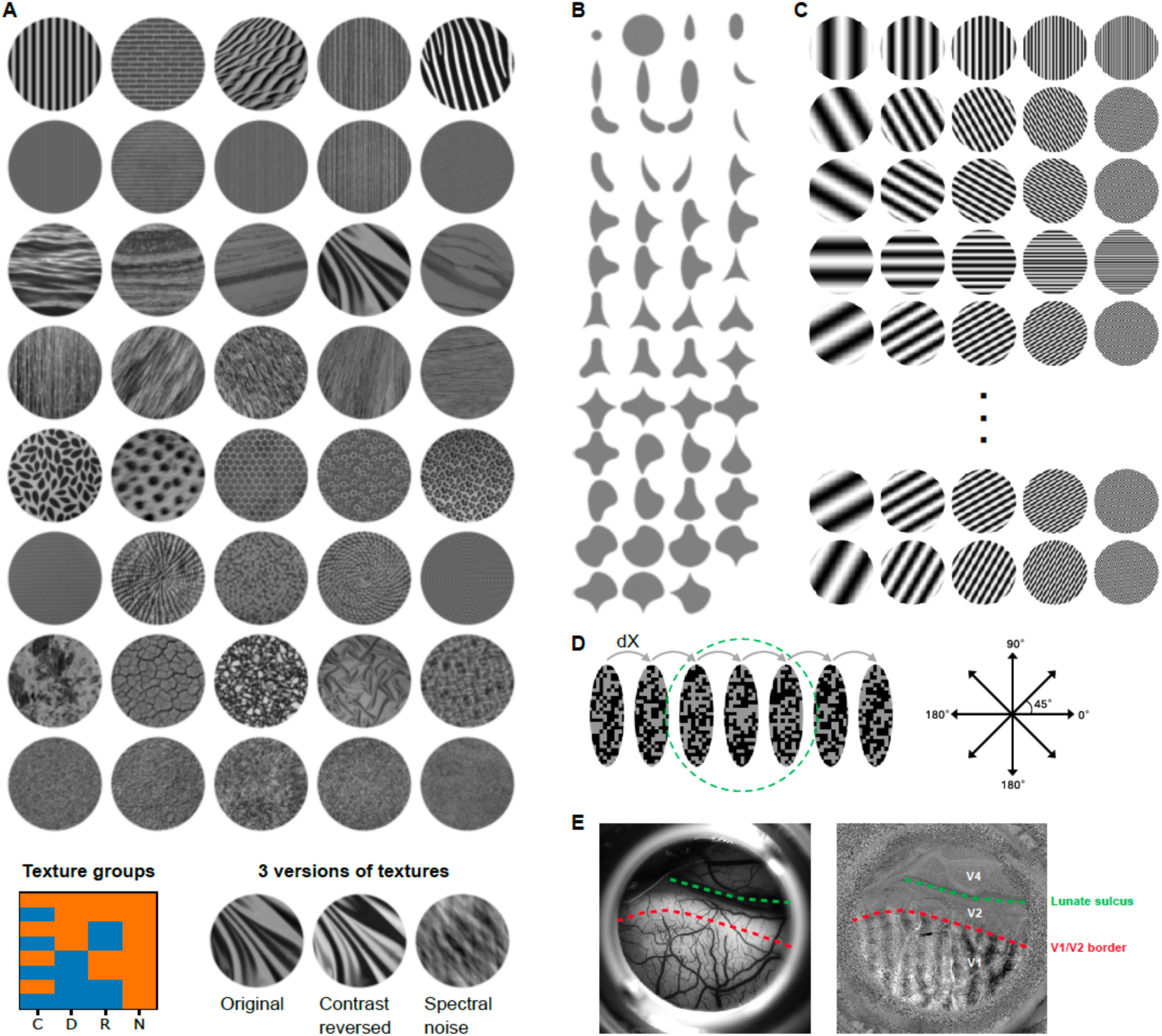
Visual stimuli and identification of the V1/V2 border. **A.** Texture stimuli. We utilized 40 naturalistic texture images categorized into eight groups (rows) based on three dimensions (C, D, R; see texture groups legend) that influence human texture perception. Each dimension takes on two levels: C: coarse (orange) versus fine (blue), D: directional (orange) versus non-directional (blue), and R: regular (orange) versus irregular (blue). For example, textures in the first row are coarse, directional, and regular, whereas those in the last row are fine, non-directional, and irregular. Each texture was shown in three variations: the original, contrast-reversed, and spectrally-matched noise. The fourth column of the texture groups legend represents the naturalness (N) dimension: the original and contrast-reversed variations are natural (orange) and spectrally-matched noise is not (blue, not-shown). **B.** Shape stimuli. One rotation of the 2D shape set from (Pasupathy and Connor, 2001) is shown. A randomly selected subset (n = 50) from rotated versions of these stimuli (n = 366, see Methods) were used to study shape responses in 17/20 sessions; in the remaining sessions (3 out of 20), a subset of 120 stimuli was used. **C.** Grating stimuli. Drifting sinusoidal gratings (4 Hz, 1 s) were presented at 12 directions of motion (30° steps) and five values of spatial frequency (0.25, 0.5, 1, 2, 4 cycles/deg). **D.** Translational motion stimuli. Elliptical noise patches with mean luminance matched to the background were displayed in sequence at seven locations centered on the receptive field (green circle). To ensure that the motion signal relied on non-Fourier cues, a unique noise pattern was randomly generated for each location on every trial. The long axis of the elliptical patch was oriented orthogonal to the direction of motion, which varied in increments of 45°. Five spatial displacement (dX) levels were tested while keeping motion speed constant (see Methods). The figure shown here corresponds to dX = 1/3 RF diameter. **E.** The exposed cortical surface (left) and imaging of ocular dominance (right) to identify V1/V2 border from one experiment. The brighter and darker stripes are right-and left-eye dominant regions in area V1, respectively. V2 recordings were made posterior to the lunate sulcus but anterior to the ocular dominance bands of V1. Note that V1-V2 segregation based on the ocular dominance pattern aligns well with a transition from denser to sparser vascular patterns from V1 to V2.

Each of the 40 texture images were presented in one of three versions, (1) the original image, (2) a contrast-reversed version in which pixel luminance was inverted, and (3) a spectrally-matched noise version, yielding a total of 120 stimulus conditions. Spectrally-matched noise images were created by removing the phase information of the original textures, by first computing the Fourier transform, randomizing the phase values, and then inverting the Fourier transform. This produces texture stimuli that retain the orientation and spatial frequency content of the original textures while removing the higher-order statistical dependencies (Freeman et al., 2013). To avoid any unintended associations between a texture category and specific local orientation energy, each texture was presented at a randomly chosen rotation on each repetition. To control for overall luminance, texture images were adjusted to have the same mean RGB pixel values (100, 100, 100). All texture images were presented through a circular aperture with a blurred boundary, and their size (8° diameter) was chosen to extend well beyond the estimated receptive field. Each stimulus was presented for a 300 ms duration, separated by 300 ms inter-stimulus interval.

#### Shape stimulus set

The shape stimulus set consisted of 50 parametric closed 2D shapes randomly selected from the original 366 shape stimulus set used to characterize V4 neurons (Pasupathy and Connor, 2001) (Figure 1B). The shapes were presented at the center of the estimated V2 RF, and their size was scaled to fit entirely within the RF boundary. Stimuli were presented for a 300 ms duration separated by 300 ms inter-stimulus interval. Since simultaneously recorded neurons may have slightly different RF locations, the shapes were presented at four additional positions shifted to the left, right, up, and down, by half of the estimated RF diameter. We then selected the optimal position for each neuron as the one that elicited the strongest mean response across all shape stimuli. In most sessions, sets of 50 unique shapes were used, but in a few sessions (3 out of 20), larger sets of 120 shapes (multiple rotations of the same shape) were used.

#### Drifting grating stimulus set

Drifting sinusoidal gratings (4 Hz temporal frequency, 1 s duration, 500 ms inter-stimulus interval) were presented at 12 directions of motion (in 30° steps) and five values of spatial frequency (0.25, 0.5, 1, 2, 4 cycles/deg) (Figure 1C). As with the texture stimuli, grating stimuli were presented through a circular aperture with a blurred boundary and were sized (8° diameter) to extend beyond the aggregate RF.

#### Translational motion stimulus set

Translational motion stimuli, like those used in (Bigelow et al., 2023), comprised of elongated noise patches presented at several sequential spatial locations centered on the estimated RF (Figure 1D). At each spatial location, the stimulus was presented for a specific duration (dT) before instantaneously moving the next location. These stimuli were designed to induce apparent motion in one of eight directions separated by 45° increments. We tested five spatial displacements (dX = 1/3, 1/6, 1/9, 1/18, or 1/36 × RF diameter), each paired with a corresponding stimulus duration (dT = 100, 50, 33.33, 16.66, or 8.33 ms) and number of spatial locations (7, 14, 21, 42, or 84), so that the motion trajectory and speed were held constant across all tested dX–dT combinations. The long axis of the elongated patches (aspect ratio = 3:1) was scaled to be 60% of the RF diameter and the total stimulus duration was 700 ms. The mean luminance of the patches was matched to the background, and a unique noise pattern was randomly generated for each of the spatial locations on every presentation; thus, stimulus motion was defined by non-Fourier cues.

### Data analysis

#### Normalized response matrix: quantification of neural response magnitude

To create a normalized response matrix for each visual stimulus set (such as the texture stimulus set), we generated response vectors for each unit. These vectors, with a length corresponding to *M* stimulus conditions (e.g., 120 texture images), were created by counting spikes within a window from stimulus onset to 100 ms after stimulus offset. We then normalized each vector by its maximum response. Finally, the normalized response vectors from simultaneously recorded neurons (ordered by channel positions along the recording probe) were aggregated to form the response matrix (*M* stimulus conditions × *N* units).

#### Quantification of similarity in feature selectivity

To quantify the similarity in feature selectivity among simultaneously recorded neurons, we computed the Pearson’s correlation coefficient between the normalized response vectors (described above) of all neuron pairs. These similarity values populated an *N* × *N* matrix, where *N* denotes the number of simultaneously recorded neurons. Values along the diagonal of this matrix, which are always 1 by definition, were replaced with NaNs; the off-diagonal values, ranging from -1.0 to 1.0, provided a measure of tuning similarity between neuron pairs.

#### Neuronal clustering analysis

We used an ad hoc clustering algorithm to identify neuronal clusters with similar feature selectivity. We first divided neurons into two groups: “in-cluster” and “out-cluster”. The “in-cluster” group consisted of neurons that exhibited statistically significant correlations with at least five of their ten nearest neighbors within the correlation matrix. Neurons that do not satisfy this criterion were categorized as “out-cluster”. After this preliminary grouping, we applied *K*-means clustering to the normalized response matrix of the “in-cluster” neurons. The *K*-means algorithm partitioned these neurons into distinct clusters based on their similarity in feature selectivity. To determine the optimal number of clusters, we used silhouette analysis, which effectively balances the need to minimize within-cluster variance while maximizing between-cluster separation. *K*-means clustering often yields a cluster with very small sizes. Therefore, as a final step, we implemented two post-processing procedures to refine the clustering results. First, we calculated the mean response vector for each cluster and merged clusters when the correlation between vectors exceeded a threshold (*r* > 0.6). Second, we removed clusters that included fewer than five neurons.

#### Orientation energy model

To determine whether a linear combination of V1-like oriented filter outputs can explain V2 shape responses, we computed the orientation energy of visual shapes using a steerable pyramid algorithm (Simoncelli and Freeman, 1995). Each shape image was decomposed using a steerable pyramid transform with four orientations and three scales, and the power of each filtered sub-band was calculated by squaring the filter responses. As orientation energy remained consistent across different scales, we focused on the power from the intermediate scale for subsequent analyses. We modeled shape responses as a linear weighted sum of four orientation filter energies. The overall significance of this model was assessed using the *F*-statistic. To quantify the similarity of orientation model fits between neurons, we computed the Pearson’s correlation coefficient of weight vectors for all neuronal pairs.

#### Direction selectivity index

For responses to drifting sinusoidal gratings and translational motion patch stimuli, we first identified orientation selective neurons by performing a one-way ANOVA with stimulus orientation as the factor. Subsequently, for neurons exhibiting significant orientation selectivity, we quantified their direction selectivity using the direction selectivity index (DSI) given by *DSI* = 1 - (*np* / *p*), where *p* represents the stronger response and *np* the weaker response along the two opposite directions orthogonal to the preferred orientation. The DSI values reported in this study are based on baseline (calculated from no stimulus trials) subtracted responses. In our analysis, direction selective neurons are a subset of orientation selective neurons. Neurons with DSI > 0.5 were classified as direction selective.

#### Orientation tuning curve fitting

Sinusoidal drifting gratings were tested with 12 motion directions (6 orientations), while translational motion patch stimuli included 8 motion directions (4 orientations). To compare orientation tuning between drifting gratings and elongated translational motion patch stimuli, we fitted the tuning curves using the *von Mises function, a periodic generalization of the Gaussian distribution*.

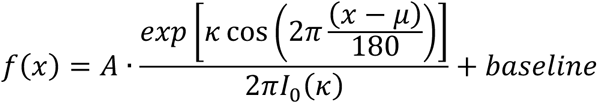

where 𝑥, 𝐴, 𝜇, 𝜅, 𝐼_0_ are stimulus orientation (in degree), amplitude, preferred orientation, tuning width parameter, and modified Bessel function of order 0, respectively.

#### Time course of texture-dependent modulation

Textures were grouped based on high and low values along each of perceptual dimension attributes (see diagrams in Figures 1 & 6), and neuronal responses to naturalistic textures (i.e., original and contrast-reversed) were compared between the groups (coarse vs. fine, directional vs. non-directional, regular vs. irregular). Additionally, we assessed the selectivity for naturalness by comparing each neuron’s mean responses to natural textures (i.e., original and contrast-reversed) versus their spectrally-matched noise counterparts.

For each single unit and stimulus condition, we extracted spike trains aligned to stimulus onset using 1 ms time bins and constructed peristimulus time histograms (PSTHs) by averaging responses across repeated trials and smoothing with a Gaussian kernel (σ = 5 ms). These PSTHs were then used to examine response dynamics over time. At each time point, we applied the Mann-Whitney U-test to assess when responses to high- and low-value stimuli along each texture dimension significantly diverged. Alternatively, some analyses relied on mean responses within defined time windows (see legends of Figure 6, 7), rather than on responses at each time point.

#### Time course of selectivity index

To assess the temporal dynamics of neuronal selectivity for texture and shape, we performed a sliding-window Receiver Operating Characteristic (ROC) curve analysis. This approach compared responses to textures grouped by high and low values along specific feature dimensions (e.g., coarseness, directionality, regularity, and naturalness), as well as to preferred (top 50%) versus non-preferred (bottom 50%) shapes. For each neuron, and at each time point (in 1 ms steps), we quantified selectivity by calculating the area under the ROC curve (AUC) based on the distributions of PSTH values for the two comparison groups—either high vs. low texture values or preferred vs. non-preferred shapes. AUC values range from 0.5 to 1, where 0.5 indicates no discriminability between conditions, values above 0.5 reflect greater responses to preferred stimuli.

#### LFP power analysis for cortical depth estimation

LFP power analysis was conducted using a 500 ms time window from –100 to +400 ms relative to visual stimulus onset. Signals were bandpass filtered using a fourth-order Butterworth filter (0.3–250 Hz) to remove slow drifts and high-frequency noise. Baseline correction was performed by subtracting the mean voltage during the 100 ms pre-stimulus period. For each channel and trial, power spectra were computed using the periodogram method from Python’s SciPy library (signal.periodogram), and then trial-averaged power was calculated. To obtain the relative power map for each probe, power values were normalized across channels at each frequency as follows (Mendoza-Halliday et al., 2024):

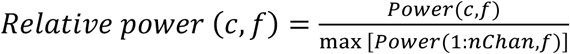

where *c, f* are channel and frequency, respectively.

From this two-dimensional matrix, we computed the average relative power within the alpha-beta (10–30 Hz) and gamma (75–150 Hz) bands as a function of channel depth. The point at which alpha-beta power minus gamma power crossed zero was interpreted as corresponding to layer 4 (Mendoza-Halliday et al., 2024). Channels with positive values were considered likely to be in superficial layers, and those with negative values in deep layers.

### Experimental design and statistical analysis

Details of experimental procedure and visual stimuli are described above. For all statistical tests presented here, independent group comparisons were performed using a nonparametric Mann–Whitney U-test, while the multiple-group comparison was done with one-way ANOVA. The statistical significance of the orientation energy model’s goodness of fit was assessed using the *F*-statistic. The strength of the linear relationship between pairs of variables was assessed by Pearson’s correlation coefficient. A *p* value less than 0.05 was considered significant.

## Results

We measured the responses of subpopulations of V2 neurons to four stimulus types— closed 2D shapes, natural textures, drifting sinusoidal gratings, and elongated object patches exhibiting translational motion (Figure 1). These stimuli have been shown to drive neurons across the V1→V2→V4 hierarchy. In particular, spatial frequency × orientation gratings have been used extensively in studies of visual neurons (Hubel and Wiesel, 1968; De Valois et al., 1982), textures tap into V2’s emerging naturalistic pattern selectivity (Freeman et al., 2013), and both closed 2D shapes and translational motion patches have revealed unique encoding properties in V4 not apparent in other early and midlevel visual areas (Pasupathy and Connor, 2001; Bigelow et al., 2023). The recordings were carried out using high-density Neuropixels probes in three anesthetized monkeys.

### Example sites: Functional clusters for shape, texture, and motion stimuli

To quantify the similarity in tuning of nearby neurons, for each stimulus type—texture, shape, drifting grating, and translational motion patches (Figure 1)—we computed the Pearson’s correlation coefficient (*r*) between the response vectors of spiking activity of all pairs of simultaneously recorded neurons. Figure 2 illustrates results from two example sites that reveal similar preference in tuning for nearby neurons across the various stimulus classes tested. In the first example (Figure 2A-D), similar responses of nearby neurons can be visualized by horizontal streaks in the response matrices (Figure 2A). For the drifting grating stimuli (the third panel, Figure 2A), two prominent horizontal bands of activity are evident in most of the simultaneously recorded neurons. Units 0 to 8 respond strongly to stimulus conditions 20 and 50 (motion direction 120° and 300°, green arrows), while units 11 to 37 show strong responses to stimulus conditions 5 and 35 (motion direction 30° and 210°, orange arrows). In both cases, the banding corresponds to two motion directions separated by 180°, indicating two sets of neurons with shared orientation selectivity. In the corresponding correlation matrix (Figure 2B), which measures tuning similarity of pairs of simultaneously recorded neurons, two clusters with high similarity in tuning (red) are evident, aligning with the neurons’ orientation tuning profiles (Figure 2A) and their spatial distribution along the probe (Figure 2C).

**Figure 2.**
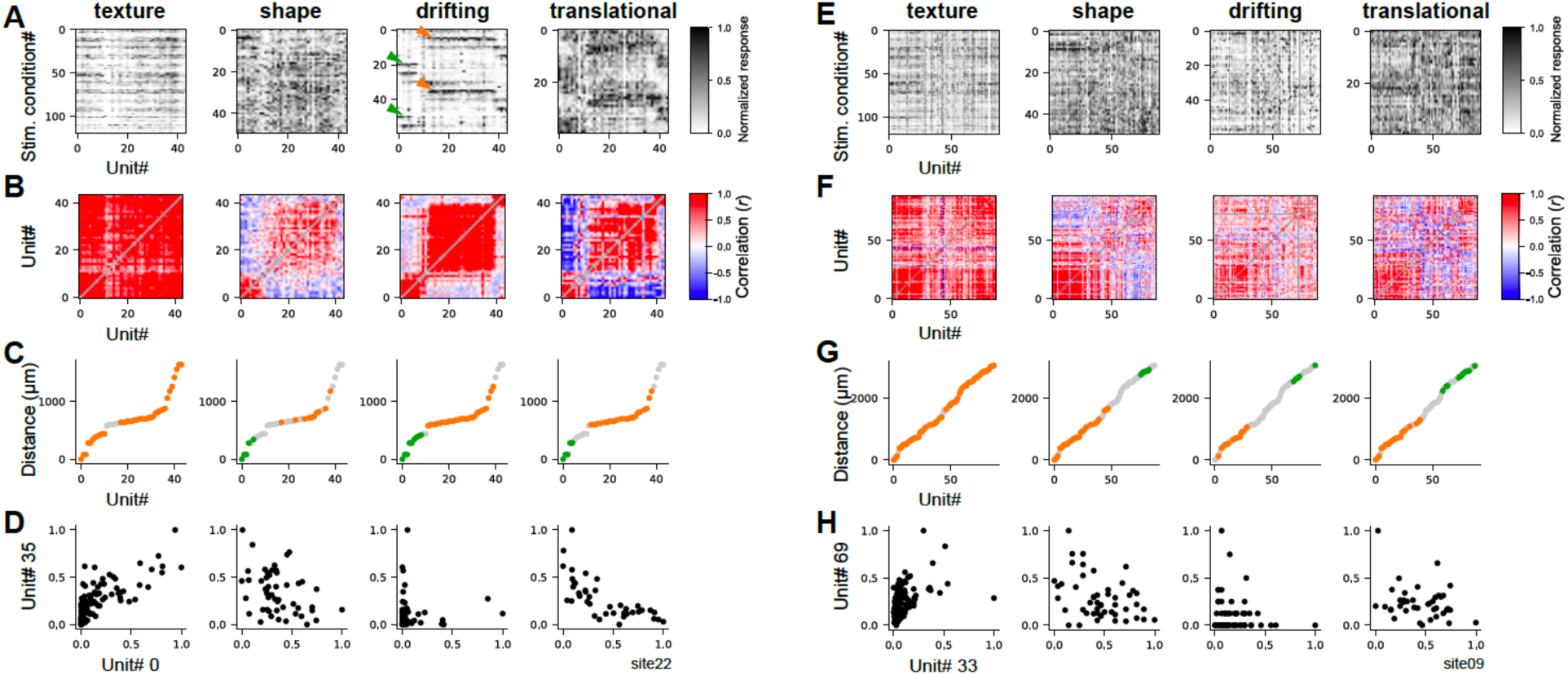
Example recording sessions. **A-D.** Data from site 22 for texture, shape, drifting grating, and translational motion stimuli (columns) are shown. **A.** Each panel shows mean normalized activity (in the 0–1 range, see scale bar) of a neuron (x-axis) across stimulus conditions (y-axis). Texture includes 120 conditions (0–39: original, 40–79: contrast-reversed, 80–119: spectrally-matched noise). Shape has 50 conditions (0–49). Drifting grating comprises 60 conditions (5 spatial frequencies × 12 directions spaced 30° apart, with 0–4 being spatial frequencies at direction 0°). Translational motion includes 40 conditions (5 spatial displacements × 8 directions spaced 45° apart, with 0–4 being displacements at direction 0°). A horizontal stripe pattern (green and orange arrows in drifting panel) indicates that multiple nearby neurons exhibit similar visual feature tuning. **B.** Pairwise tuning similarity between neurons is shown, with unit IDs on both axes. Color scale indicates the strength and sign of similarity, where red and blue represent positive and negative values for the Pearson’s correlation coefficient (*r*). Diagonal entries were replaced with NaNs and displayed in gray. **C.** The x-axis represents unit IDs, and the y-axis values correspond to the distance from unit# 0, defined as the neuron recorded closest to the probe tip. Neurons with similar tuning were organized into groups using clustering analysis (see Methods). The four panels display the same neurons, but colors represent different functional clusters for each stimulus type: orange marks the largest cluster. Neurons that do not belong to any cluster are represented in gray. **D.** Scatter plots comparing responses of two example units. These two units belonged to the same cluster and exhibited a positive correlation for texture responses but not for other stimulus classes. **E-H.** Results from site 09. The conventions are the same as those in **A-D.**

We observed similarly clustered tuning preferences for shape and translational motion patch stimuli (the second and fourth panels in Figure 2B). With texture stimuli, we observed a notably different profile: tuning similarity was consistently high across the entire length of the probe even for neurons #0 and #43 separated by 1.6 mm (the first panel, Figure 2B). For each stimulus type, we used an ad hoc clustering algorithm to group neurons into clusters based on their tuning similarity (Figure 2C; also see Methods). In the case of texture stimuli, nearly all cells exhibited similar tuning and were classified as a single group. In contrast, for shape, drifting grating, and translational motion stimuli, at least two distinct groups emerged with different tuning characteristics. Figure 2D illustrates the responses of two example units (#0 vs. #35) that belong to the same group for texture selectivity but assigned to different groups for other stimulus types; their responses are positively correlated for texture stimuli but negatively correlated for all other stimulus types. Figure 2E-H shows similar overall trends from another recording site. Together, these findings suggest that in V2, neurons exhibit clustered selectivity for a variety of stimulus classes, with texture stimuli leading to a more widespread, coordinated response across neurons.

### Population results

We quantified the prevalence of functional clusters for different visual stimulus types using data from three macaque monkeys: 24 sites (1489 neurons) for texture, 20 sites (1138 neurons) for shape, 7 sites (600 neurons) for drifting grating, and 12 sites (802 neurons) for translational motion patch stimuli. At the population level, we observed functional clusters in response to all stimulus classes; however, as with the examples in Figure 2, texture stimuli consistently evoked more extensive clustering than other stimulus types (Figure 3A). To explore this further, we examined how response similarity varies as a function of inter-neuron distance for each stimulus type (Figure 3B). For texture responses, mean similarity between neurons decreased gradually with increasing distance, remaining substantially high (mean *r* = 0.19) even at 2 mm. In contrast, for shape, drifting grating, and translational motion patch stimuli, similarity declined more sharply up to distances of 500 μm. At neuronal distances in the 1–2 mm range, similarity was generally low, with mean values of 0.02 for shape, 0.08 for drifting grating, and 0.06 for translational motion patch.

**Figure 3.**
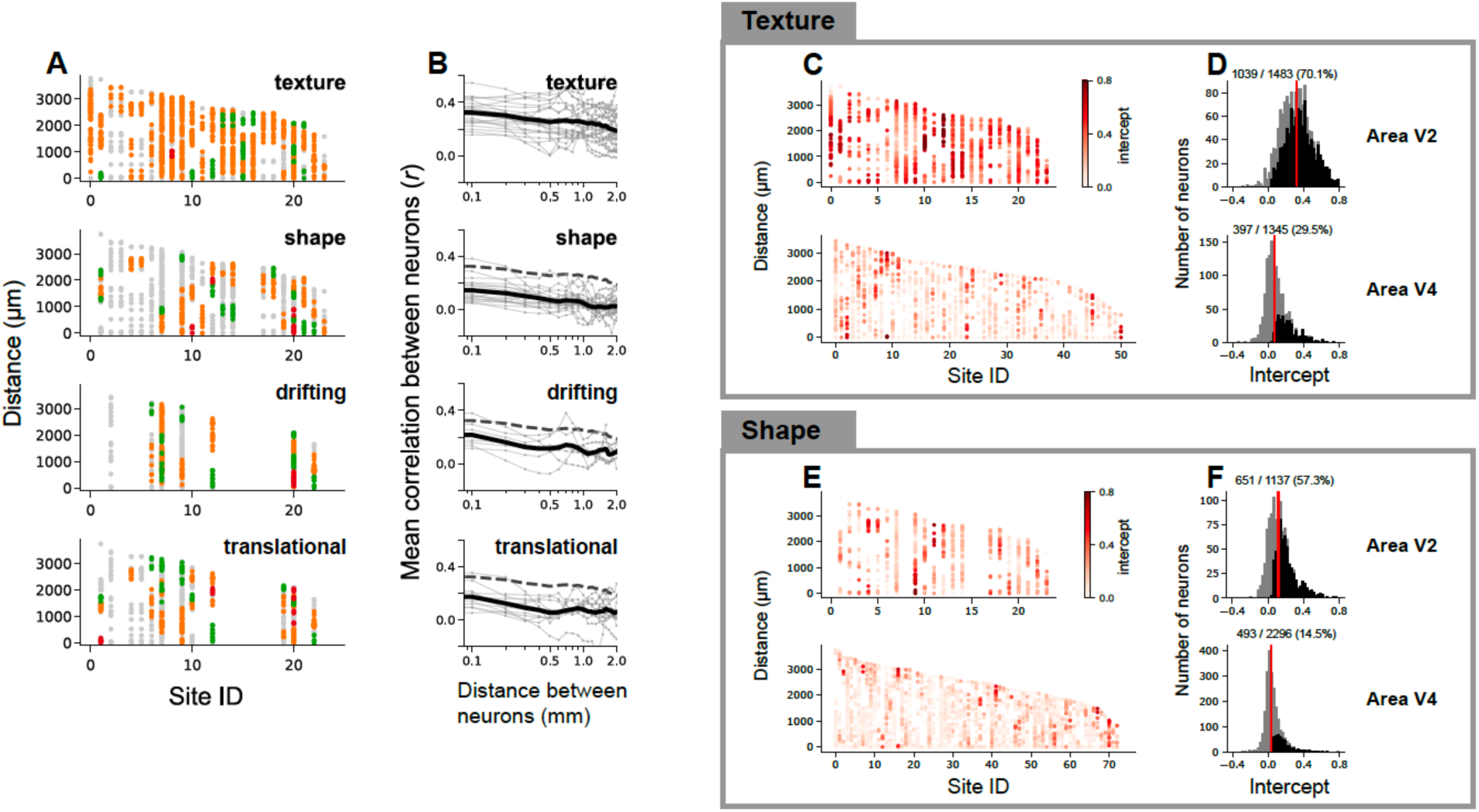
Population data comparison from V2 and V4. **A-B.** Neuronal clusters in V2. **A.** Results from clustering analysis (see Methods) across 24 recording sites (x-axis) for the four stimulus classes. In each recording site, individual neurons are represented by dots positioned at the depths of the corresponding contact along the probe. A depth of 0 corresponds to the location of the deepest neuron recorded in each session. Distinct clusters of neurons with similar tuning are color-coded, while neurons not assigned to any cluster appear in gray. Orange indicates the largest cluster within each site, followed by green for the second largest, and red for the smallest. Note that colors are meaningful only within a recording site; the same colors across different sites do not indicate shared feature selectivity. **B.** Dependence of tuning similarity on inter-neuronal distance. In each panel, thin gray lines correspond to individual recording sites, with each data point representing the mean value of tuning similarity between pairs of neurons from 300 μm bins positioned every 100 μm. The black line indicates the average value across recording sites. Mean tuning similarity for texture stimuli is replotted in the other panels (dashed line) for comparison. **C-D.** Comparison between V2 and V4. As in (Namima et al., 2025), we used a linear regression model to relate texture tuning similarity and inter-neuronal distance (see the main text for details). A high similarity among nearby neurons that decreases with distance would yield a positive intercept and a negative slope. Intercept values (color) from individual V2 (top) and V4 (bottom) neurons across all recording sites are plotted as a function of cortical depth. Many neurons in V2 but not V4 exhibit a significant positive intercept indicative of tuning similarity of neighbors. The same color scale is used for both top and bottom. **D.** Histogram of intercept values from the texture tuning similarity regressions in area V2 and V4. Black bars indicate neurons with statistically significant positive intercepts. The median value for V2 (0.32, indicated by the red line) is significantly higher than that for V4 (0.07) (Mann-Whitney U-test, p < 0.001). **E-F.** Same comparison analyses applied to shape tuning similarity in area V2 (top) and area V4 (bottom). The median value for V2 (0.12) is significantly higher than that for V4 (0.03) (Mann-Whitney U-test, p < 0.001). Note that for both shape and texture stimulus types, tuning similarity of neighboring neurons in V2 is more spatially extensive than in V4, as reflected by higher intercepts. In **C-F**, panels for V4 data were adapted from (Namima et al., 2025).

One possible explanation for the larger clusters observed in response to textures is that some of the texture stimuli may not have driven the neural activity strongly; this could induce uniformly low responses for a subset of stimuli across the population thereby inflating tuning similarity. In our recordings, most neurons were indeed more strongly modulated by coarse textures (associated with lower spatial frequencies) than by fine textures, and by natural textures rather than noise textures (see *Neuronal clusters encoding different perceptual texture dimensions*). To address this potential explanation, we performed two analyses. First, we asked whether larger neuronal clusters for texture were observable even when responses to fine or noise textures were excluded from the response similarity calculations. Figure S1A-C shows this is indeed the case. In addition to analyses focused on texture stimuli, comparative analyses involving shape stimuli were also conducted, confirming that driven firing rates and response variability were similar between texture and shape stimuli (see Figure S1D-E). Second, we assessed response similarity between random pairs of neurons across different recording sites, reasoning that if response similarity is inherent to our stimulus design, the strong similarity will remain between responses of randomly paired neurons. Across 10,000 random pairs, the mean similarity was 0.16 ± 0.002 (mean ± SEM)—closely matching the values observed between neurons separated by 2 mm within the same recording site. These findings suggest that the observed clustering for texture stimuli reflects genuine functional organization and is not simply an artifact of limited stimulus drive by a subset of stimuli.

Our results in V2 provide a striking contrast to the sparse functional clustering we recently observed in V4 with the same shape and texture stimuli used here (Namima et al., 2025). To directly compare the results from V4 and V2, we applied the same analytical approach used in our previous V4 study to our V2 data. Specifically, for each neuron, we fitted a linear regression model to capture how response similarity between the neuron in question and all other neurons within the recording site varied as a function of inter-neuronal distance. When a neuron is part of a functional cluster, this relationship is typically well fit by a model with a positive intercept and a negative slope. When we examined the proportion of V2 neurons with statistically significant positive intercept parameters (p < 0.05), we found that approximately 70% of neurons met this criterion for texture stimuli and 57% for shape stimuli (Figure 3C-D, E-F). These proportions are substantially higher than those reported in area V4, where only about 20% of neurons exhibit tuning similar to their neighbors, suggesting a substantial change in functional organization between V2 and V4.

### Dissecting the representational bases of clustered encoding in V2

Prior studies in V2 have demonstrated selectivity for basic visual features such as local orientation, spatial frequency, and motion direction (Foster et al., 1985; Levitt et al., 1994; Gegenfurtner et al., 1996; Felleman et al., 2015), as well as a preference for naturalistic textures over spectrally-matched noise (Freeman et al., 2013; Ziemba et al., 2016; Okazawa et al., 2017). Optical imaging has further revealed columnar organization for orientation and motion direction (Lu et al., 2010; Hu et al., 2018). Here, we examine whether the selectivity and spatial clustering observed in response to our more complex stimuli can be accounted for by tuning and clustering based on lower-level visual features. Specifically: (1) we asked how orientation energy contributes to the clustering of shape responses, (2) we compared direction and orientation selectivity for drifting versus translational motion, and (3) we characterized texture selectivity of neuronal clusters in terms of human perceptual dimensions.

#### Clustering of shape responses in V2 can be explained by a weighted sum of orientation energy

Area V2 is known to encode combinations of orientations (Ito and Komatsu, 2004; Anzai et al., 2007), potentially supporting the representation of more complex visual features such as angles, contours, or textures. We sought to investigate whether the shape tuning of V2 neurons, and their clustered shape preferences, could be described in terms of weighted combinations of V1-like oriented filter outputs.

For each neuron, we determined the weights for four orientation-tuned filters to best fit the shape responses using linear regression (see Methods). Our analysis showed that the orientation energy model provided a statistically significant fit to the shape responses in approximately half of the V2 neurons tested with shape stimuli (517 out of 1138 neurons, 45.4%) (Figure 4A-B). Furthermore, we observed a strong correspondence between similarity matrices derived from orientation weight vectors (four filters) and those based on the full set of shape responses (50 shapes) (Figure 4C). This finding suggests that for a substantial portion of V2 neurons, shape selectivity and clustering based on shape responses can be largely explained by a linear combination of V1-like oriented filter outputs. When the same analysis was applied to V4, approximately 35% of neurons (804 out of 2,295) exhibited a statistically significant fit, with the overall explained variance being significantly lower than in V2 (Mann-Whitney U-test, p < 0.001). However, since the number of shapes (n = 50) used was limited, a more precise comparison between the two areas would require a larger and more diverse set of stimuli to fully capture the differences in shape selectivity and underlying neural coding mechanisms (Oleskiw et al., 2014).

**Figure 4.**
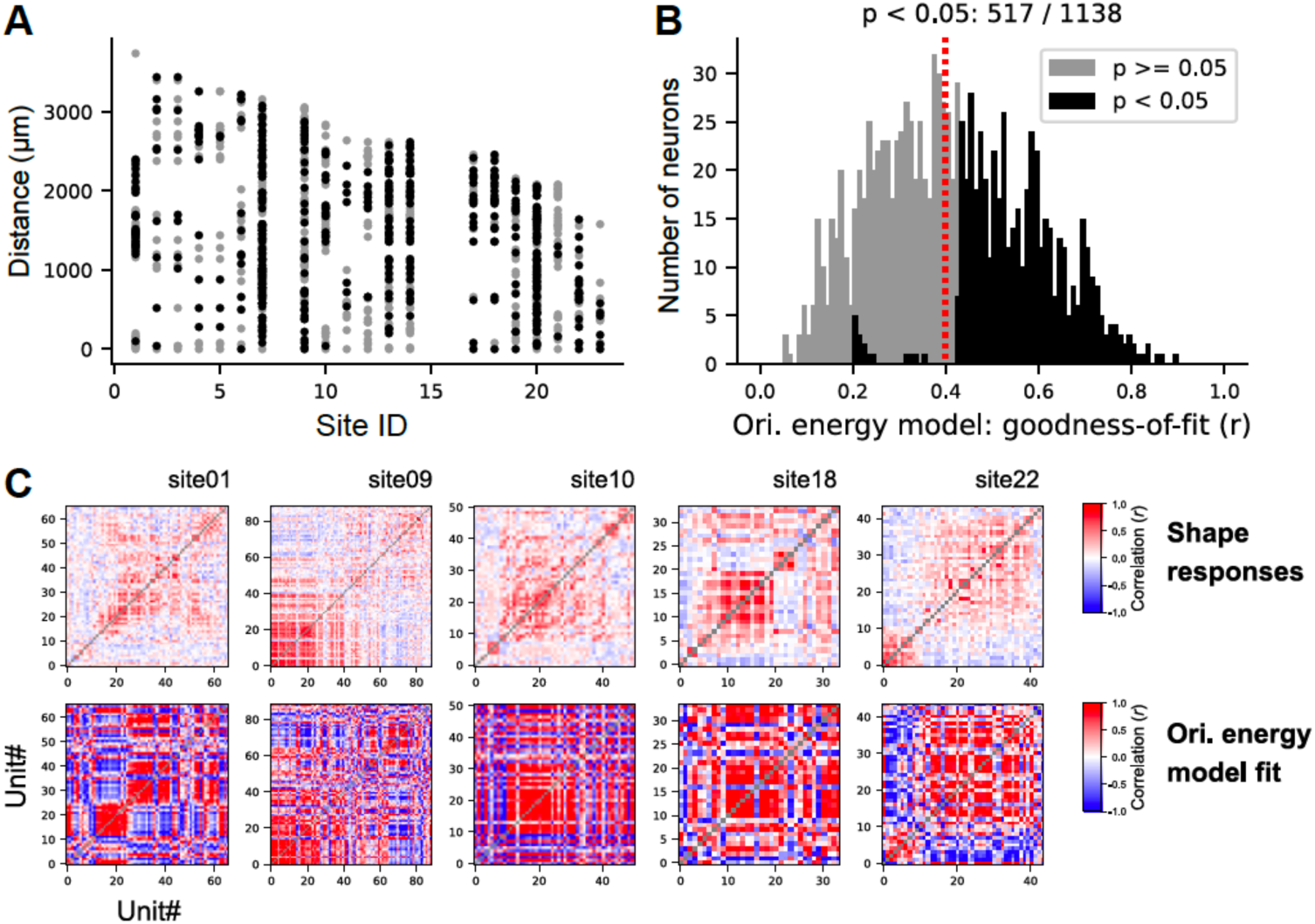
Orientation energy model fit to shape responses. **A-B.** For each neuron, shape responses were predicted using an orientation energy model based on linear regression. Black dots in **A** and black bars in **B** represent neurons with statistically significant goodness-of-fit (*r*) as determined by an *F*-test (p < 0.05), while grey dots and bars represent neurons that did not reach statistical significance. In most sessions, we used a shape set of 50 stimuli, but in a few sessions (3 out of 20), a larger set (n=120) was used, which led to statistical significance even with smaller correlation values (*r* < 0.4). The vertical line represents the median (0.40). **C.** A comparison of the tuning similarity matrices derived from measured (top) versus predicted (bottom) responses from the orientation energy model. Matrices exhibit qualitatively similar structure.

#### Clustering of responses to motion stimuli: contributions of surface and object-based motion computations

Previous studies have predominantly used drifting gratings and fields of dynamic dots to probe motion processing in visual cortex (Britten et al., 1992; Snowden et al., 1992; Livingstone, 1998; Gur et al., 2005). These surface motion stimuli, where a visual pattern drifts within a static aperture, have been valuable for measuring the selectivity and clustering of direction selective neurons in V1, V2 and MT (Albright et al., 1984; Hawken et al., 1988; Lu et al., 2010). Recently, using translational motion stimuli, we identified a sub-group of neurons in V4 that encode motion direction only when defined by the displacement of a stimulus patch across the RF. These “object motion” sensitive neurons encode motion over a large range of spatial displacements even when defined by non-Fourier cues (Bigelow et al., 2023) unlike neurons in the quintessential motion area MT (Churchland et al., 2005; Hedges et al., 2011). Motivated by these findings, we used the same stimuli (Figure 1C-D) to examine the clustering of motion responses in V2 and identify whether area V2 shows distinctive sensitivities for drifting and translational motion.

Across our dataset, ∼44% (179 out of 408) of orientation-tuned V2 neurons were direction selective in response to drifting motion stimuli, and a smaller proportion (∼29%; 141 out of 486) were direction selective for translational motion stimuli for the largest displacement level (dX = RF/3) we tested (Figure 5A-D). For both stimulus classes, direction selective neurons were not randomly distributed but exhibited a clustered organization within specific recording sites (e.g., sites 7, 9, 20; Figure 5A, C), suggesting localized cortical domains of direction selectivity. This clustering occurred even though drifting and translating stimuli were defined by Fourier and non-Fourier cues, respectively. Overall, clustering was more prominent with drifting that with translational motion, but in both cases, neurons tuned to two opposite directions (i.e., 180° apart) were intermixed (Figure 5I). For a given neuron, direction tuning curves with drifting and translational motion stimuli were similar in overall shape, although the peak and strength often differed. For example, in Figure 5E-H, the neuron shown in blue exhibited stronger direction selectivity with drifting motion stimuli; the neuron in red preferred opposite directions (90° and 270°) when tested with drifting versus translational motion stimuli. Across the population, motion direction selectivity was stronger for drifting than translational motion (Figure 5I, J) and the difference in peak showed a bimodal distribution, i.e., tuning for drifting and translational motion were in the same or opposite directions (Figure 5K).

**Figure 5.**
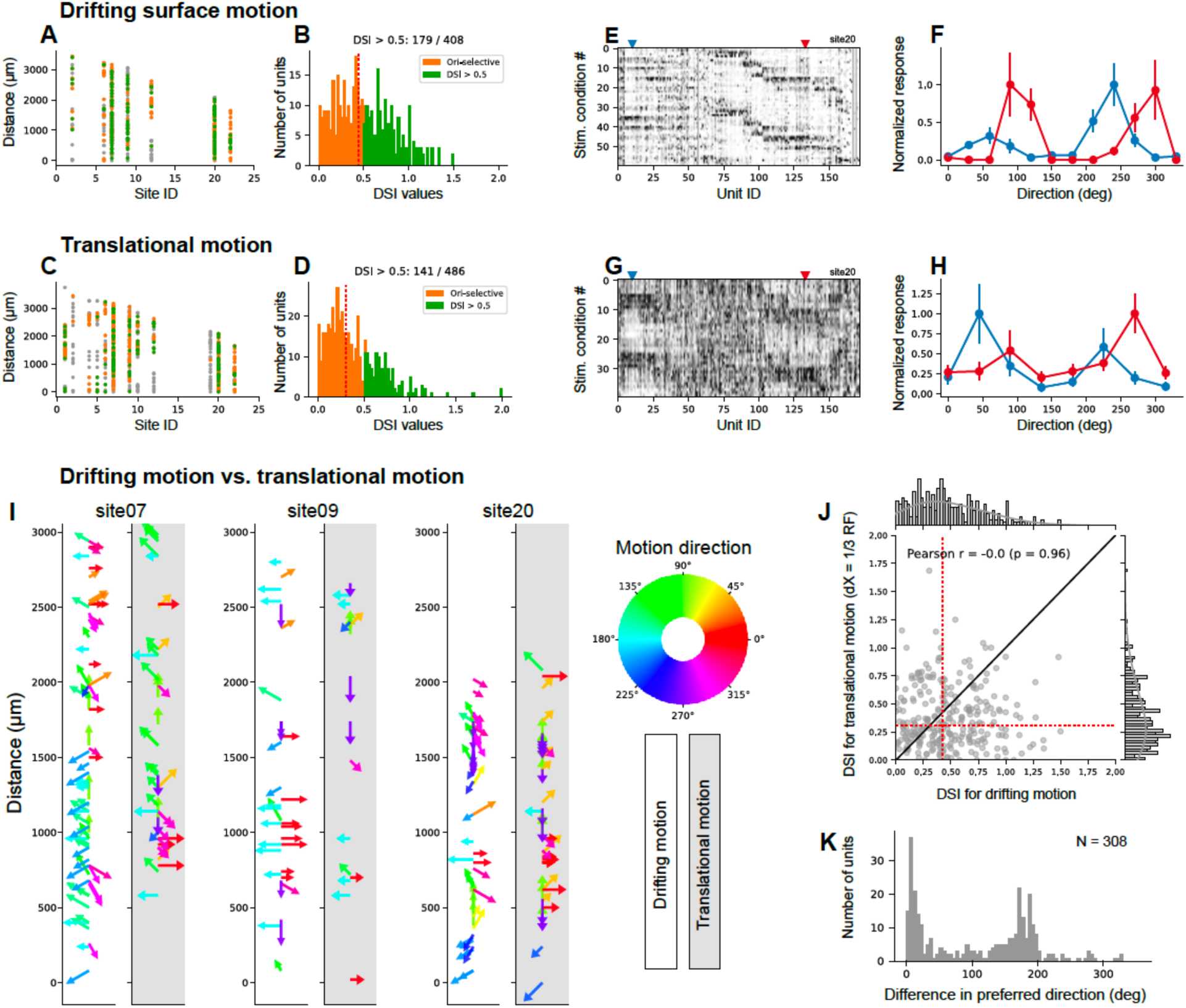
Comparing direction selectivity for drifting and translational motion. **A.** Individual neurons from seven recording sites tested with drifting motion stimuli are shown as dots and colors denote orientation selective neurons (orange), direction selective neurons (green) and neither (gray). **B.** Histogram of the direction selective index (DSI) for orientation and direction selective neurons (using the same color scheme as **A**) computed at the optimal spatial frequency determined for each neuron. **C-D.** Results from 12 recording sites (the seven sites shown in **A** plus five others) tested with translational motion stimuli are shown using the same convention as **A-B.** DSI values were computed at the largest spatial displacement (dX = RF/3). **E.** Normalized response matrix for drifting grating stimuli (60 conditions = 12 directions × 5 spatial frequencies) from one session (site 20), following the same convention as in Figure 2A. A horizontal stripe pattern indicates that multiple nearby neurons exhibit similar visual feature tuning. **F.** Direction tuning curves from two example neurons from the session in **E**. The corresponding neurons are marked with the same colors in panel **E**. Error bars represent the standard error of the mean. **G.** Normalized response matrix for translational motion stimuli for the same session in panel **E. H.** Tuning curves for translational motion stimuli for the same two neurons in **F**. Red/blue tuning curve shapes are similar (compare panels **F** and **H**), but direction selectivity for drifting motion differs from that for translational motion. **I.** The optimal direction (color) and DSI value (arrow length) for each stimulus type and each neuron from three recording sites (sites 07, 09, 20) are shown. Clustered preferences for direction and orientation can be observed. **J.** DSI for drifting grating stimuli (x-axis) and translational motion stimuli (y-axis). The data revealed no correlation between the two measures (*r* = -0.02, p=0.96). Red dotted lines indicate median DSI values for translational (0.304) and drifting (0.423) motion, respectively. **K.** Histogram of peak difference between drifting and translational motion tuning curves across 308 neurons that exhibited orientation selectivity for both stimulus types. The histogram reveals a bimodal distribution, indicating that the difference in preferred direction is typically either near 0° or 180°.

These results suggest that direction selectivity for drifting and translational motion may arise from differential modulation of an underlying orientation tuning curve. Unlike V4, we found no evidence in V2 for a distinct object-based motion processing system that is sensitive to translational but not drifting motion (Figure S2A-B; (Bigelow et al., 2023)). V2 is also unlike MT, which is insensitive to non-Fourier motion cues (O’Keefe and Movshon, 1998) and exhibits a loss of direction selectivity for translational motion when spatial displacement is large (Figure S2C-D; (Mikami et al., 1986; Churchland et al., 2005)). Overall, our results support the hypothesis that V2 exhibits clustered selectivity for surface motion defined by Fourier and non-Fourier cues over a broad range of spatial displacements.

#### Neuronal clusters encoding different perceptual texture dimensions

Previous studies have demonstrated that responses of V2 neurons can be modeled in terms of higher-order texture statistics (Okazawa et al., 2017), but it remains unclear whether this selectivity can be explained in terms of tuning to specific perceptually defined dimensions of texture as observed in V4 (Kim et al., 2022) and whether spatial clustering exists for these dimensions. To address this, here we examined the selectivity of individual neurons along four dimensions—the perceptual dimensions of coarseness, directionality, and regularity (Kim et al., 2022), and a naturalness metric (see Methods)—and asked how this preference compared across neurons within a recording site. Results from one recording site are illustrated in Figure 6A (the same recording site as Figure 2A). Across recorded neurons, strong responses represented by dark horizontal strips in the response matrix (Figure 6A, left) mirror the stimulus grouping along the coarseness dimension (Figure 6A, right). Responses were also stronger to both the original (top) and contrast-reversed (middle) naturalistic textures than to spectrally-matched noise counterparts (bottom). Temporal dynamics (Figure 6B) also reveal a preference for coarse, irregular and natural textures across most neurons in this recording site.

**Figure 6.**
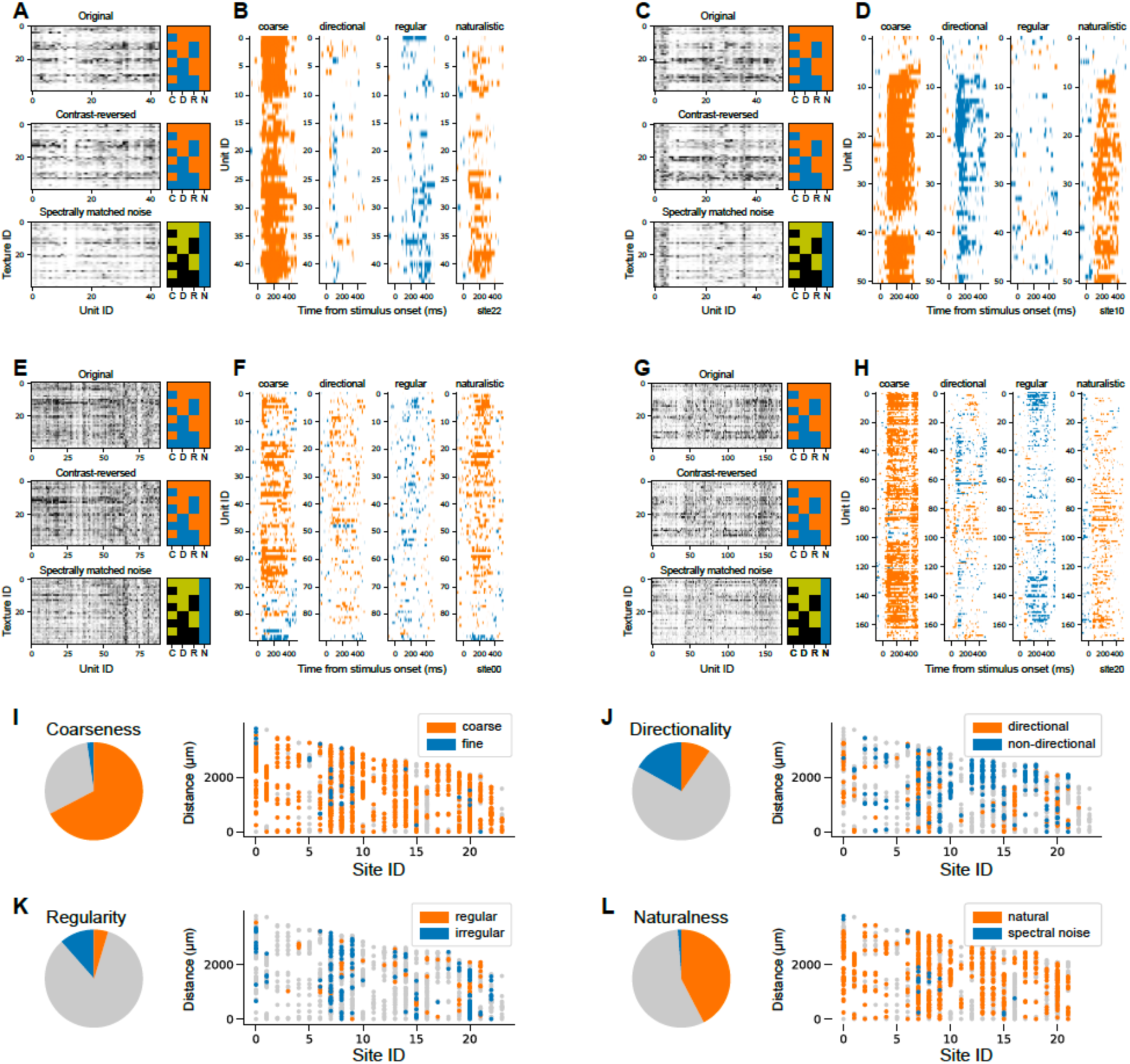
Neuronal clusters with different texture selectivity. **A.** Normalized response matrix (x-axis: unit ID, y-axis: texture ID) for an example session and the temporal response profile along the four texture dimensions (right). Responses to three texture variations are shown: original (top), contrast-reversed (middle), and spectral noise (bottom). The texture dimensions, C, D, and R follow definitions in Figure 1. The naturalness dimension (N) takes on two values: original and contrast-reversed are natural (orange) while spectral noise is not (blue). In the spectral noise condition, C, D, and R dimensions are marked in green and black rather than orange and blue, since they were not included in the computations shown in **B. B.** For each neuron, time points with statistically significant differences in mean responses between the two levels along each texture dimension (e.g., coarse vs. fine) were identified and marked as orange or blue corresponding to the greater value (see Methods). For C, D, R dimensions, only responses to naturalistic textures were used in comparisons. Note that nearly all neurons exhibit similar texture selectivity associated with a preference for coarse, irregular, and naturalistic features. **C-D.** The same analyses as in **A-B** were performed for site 10. Nearly all neurons demonstrate similar texture selectivity, showing a preference for coarse, non-directional, and naturalistic features. **E-F.** Results from site 00. Nearly all neurons demonstrate similar texture selectivity, showing a preference for coarse, directional, and naturalistic features. **G-H.** Results from site 20. Clusters of neurons with slightly different preferences for directionality and regularity were identified across different depths, but overall texture selectivity remained similar. **I.** Across all recording sites, the proportions of neurons significantly modulated by coarseness (left) were calculated from mean responses within the 0–400 ms post-stimulus window, and their positions were mapped along the probe (right). Grey dots indicate neurons with no preference. **J-L.** The same analyses as in **I** for directionality (**J**), regularity (**K**), and naturalness (**L**) features.

A preference for coarse and naturalistic texture features was commonly observed across recording sites (Figure 6I, L). However, selectivity for other texture attributes varied substantially across individual neurons and sites. For example, neurons recorded at site 10 showed a preference for coarse, non-directional, and naturalistic textures with minimal influence from regularity (Figure 6C-D). Data from other example sites are shown in Figure 6E-F, G-H, further demonstrating that V2 contains specialized subpopulations tuned to different texture attributes, while maintaining a general bias toward naturalistic features (for more detailed examples of single unit activity, see Figure S3).

### Distinct temporal dynamic patterns are differently distributed across cortical layers

In previous sections, our analyses of functional clustering in V2 neurons were based on time-averaged responses without considering their temporal dynamics. To investigate whether incorporating temporal dynamics reveals additional structure, we analyzed the evolution of feature selectivity over time and assessed whether distinct temporal dynamics are systematically organized across layers.

As a first step, we clustered neurons based on the similarity of their temporal response profiles (PSTHs). Our analysis used a *K*-means clustering approach and was constrained to a 300 ms window (−100 ms to 200 ms) around stimulus onset. This identified three clusters with distinct response dynamics. In Figure 7A, neurons in the black cluster exhibited a short-latency (∼65 ms) transient peak that rapidly decayed. In contrast, neurons in the blue cluster showed a slightly delayed peak at ∼90 ms, and rather than decaying quickly, responses were maintained as sustained activity persisting well beyond 100 ms after stimulus onset. Lastly, neurons in the green cluster showed a slower rise in activity without a distinct peak, but responses converged with those of the blue cluster after 130 ms.

**Figure 7.**
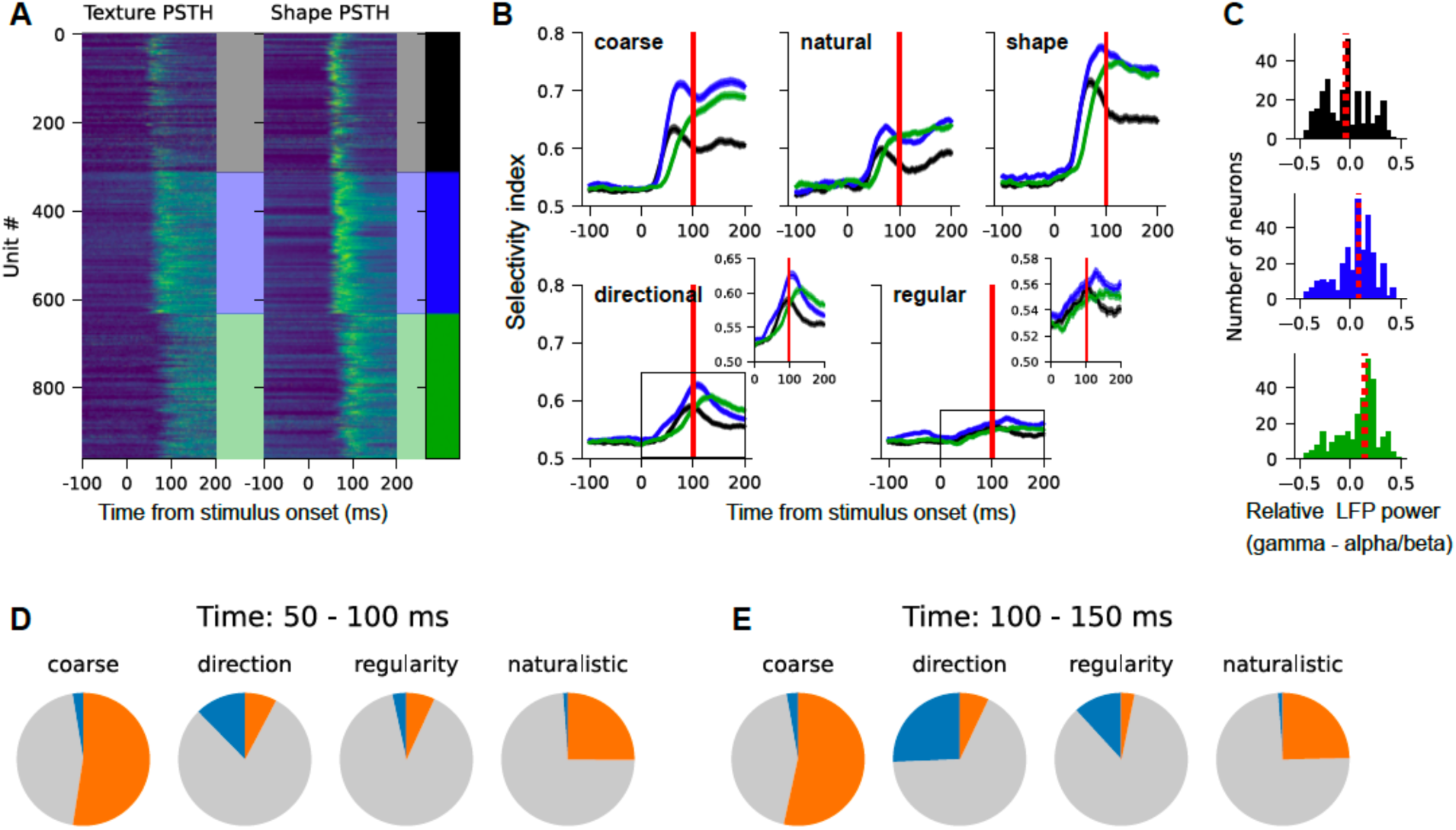
Temporal dynamics of texture selectivity and its relationship with cortical depths. **A.** All recorded V2 neurons with clear visual responses were grouped into three clusters using *K*-means algorithm, based on the similarity of their temporal dynamics from –100 to 200 ms relative to stimulus onset. The PSTHs in the black cluster are characterized by a fast transient peak, whereas those in the blue and green clusters exhibit stronger sustained activity, persisting 100 ms after stimulus onset. **B.** For each PSTH cluster, selectivity indices (i.e., area under the ROC curve; see Methods) for four texture dimensions and shape are plotted as a function of time. Selectivity for coarse, naturalistic texture features and shape emerged earlier than that for directional and regular texture features. Notably, neurons with sustained responses (blue and green clusters) exhibited stronger overall selectivity. **C.** For each PSTH cluster, cortical layers of individual neurons were estimated by the difference of relative LFP power between gamma (50–150 Hz) and alpha–beta (10–30 Hz) frequency ranges. Positive and negative values indicate neurons located in the superficial and deep layers, respectively, with values near 0 corresponding to layer 4. **D-E.** The relative proportions of neurons with statistically significant modulation for each texture attribute. Two distinct time windows were analyzed to contrast early (**D.** 50-100 ms post-stimulus onset) and later (**E.** 100-150 ms post-stimulus onset) neuronal activity. Neurons with significant modulation are represented by orange and blue colors, with color schemes consistent with those in Figure 6.

Importantly, the temporal dynamics of individual neurons within each cluster were highly consistent across texture and shape PSTHs (left vs. right in Figure 7A), although peak responses were stronger for shape. Neurons exhibiting sustained activity patterns (blue, green) demonstrated significantly stronger selectivity for both shape and all texture features compared to those with transient responses (black) (Figure 7B), suggesting that these neurons may play an important role in supporting both texture and shape processing in V2.

To assess whether these neuronal groups also differed in their laminar distribution, we estimated the laminar positions of individual neurons based on the relative differences in local field potential (LFP) power spectra between the alpha/beta and gamma frequency ranges (Mendoza-Halliday et al., 2024) (see Methods & Figure S4). While both sustained and transient neurons were distributed across cortical layers, sustained neurons were more frequently found in superficial layers (Figure 7C). In contrast, transient neurons were slightly more common around layer 4, suggesting a possible association with initial feedforward processing. These findings suggest that texture and shape processing in V2 is organized not only in time but also across cortical depth, with distinct layers contributing preferentially to early versus sustained encoding of visual features.

Our recent V4 study found that response modulation related to directionality and regularity are significantly delayed than those for coarseness and contrast (Kim et al., 2022). To determine whether similar results are observed in V2, we quantified the proportion of neurons exhibiting statistically significant modulation for each texture attribute within two post-stimulus time windows: early (50–100 ms) and late (100–150 ms) phases (Figure 7D-E). In the early phase, a large proportion of neurons were modulated by coarseness and naturalistic features (large orange slices), while fewer neurons responded to directionality and regularity. In the later phase, the proportion of neurons responding to coarseness and naturalistic features remained largely unchanged, but there was a marked increase in neurons sensitive to directionality and regularity (indicated by the larger blue slices). This analysis showed that selectivity for coarse and naturalistic features emerges rapidly, whereas sensitivity to directionality and regularity develops more gradually during the response. We did not observe a clear relationship between these response dynamics and cortical layers.

In summary, V2 neurons form large clusters that jointly encode multiple perceptual texture dimensions. Both texture and shape selectivity emerge gradually across time and cortical space, with sustained dynamics in superficial layers further refining the representation.

## Discussion

We used high-density Neuropixels probes to study populations of single neurons and determine the functional organization and the representational bases of area V2. Our results provide the first documentation of clustered selectivity across cortical layers in V2 for higher-order form, texture, and motion stimuli. The organization we observed rivals those previously demonstrated in V1 and is strikingly different from the sparse clusters observed in V4 with similar stimuli (Namima et al., 2025). V2 clusters based on responses to texture stimuli were spatially more extensive than those to other stimuli, suggesting that texture information may be integrated over broader cortical networks than those processing spatially localized features such as contours or object boundaries. In terms of representational bases, V2 encodes visual stimuli largely in terms of surface characteristics, supporting the hypothesis that object-based coding emerges first in V4 (Pasupathy et al., 2020). Overall, these results provide new insights into the representational transformations across the early and midlevel stages of visual processing, offering constraints to refine future hierarchical models of the ventral stream.

### Functional organization: relationship to prior studies

V2 exhibits a well-defined modular organization characterized by repeating patterns of thin, thick, and pale stripes, identifiable histologically through cytochrome oxidase (CO) staining (Horton, 1984). Functional maps from optical imaging have demonstrated the existence of distinct functional clusters for features (e.g., orientation, color, motion) corresponding to these stripes (Ts’o et al., 1990; Lu and Roe, 2008; Felleman et al., 2015).

These approaches focused on properties across the cortical surface and neuronal subpopulations at a coarse level (although a few studies achieved finer resolution; see (Hubel and Livingstone, 1985; Shipp and Zeki, 2002)). Our study expands this understanding by demonstrating robust clustering at the single-neuron scale across cortical layers for higher-order stimulus dimensions.

With regards to the size of clusters, we found that similarly tuned neurons spanned ∼500 μm along the length of the probe for shape, drifting grating and translational motion patch stimuli. We may not have seen larger clusters encompassing all cortical layers, because our probes were often inserted at an oblique angle traversing multiple orientation domains (see Figure 2, 5). The number of columns crossed was variable across penetrations and dependent on the penetration angle (ranging from perpendicular to parallel to the cortical layers). With texture stimuli, we observed much larger functional clusters, with response similarity decreasing more gradually and remaining relatively high even beyond 1 mm. The neural basis for this difference is unclear, but we posit that these larger clusters for texture stimuli may arise because texture selectivity is built by integrating higher-order correlations of stimulus features across space. Indeed, orthogonal coding of orientation and spatial frequency has been reported (Zhang et al., 2023). The texture clusters we identified, which prefer coarse features and extend across multiple orientation columns, may in part reflect the fact that neuronal groups tuned to low spatial frequencies necessarily encompass neurons with diverse orientation preferences. However, weaker responses to spectrally-matched noise textures with identical spatial frequency content, and varied preference for higher-order features (e.g., directionality, regularity) across recording sites indicate that spatial frequency alone is insufficient to explain these effects.

Our current results are strikingly different from our previous work in V4, where neurons tuned to specific shapes and textures formed very sparse clusters; clearer clustering was observable only at a coarser stimulus category level (e.g., horizontal versus vertical orientations) and at the population level, as revealed with optical imaging (Tanigawa et al., 2010; Li et al., 2013) or by pooling activity over spatial scales of hundreds of microns (Namima et al., 2025). Thus, our results highlight a dramatic shift in modular structure from V2 to V4.

### Columnar organization in the context of area size and feature complexity

Our findings are pertinent to the longstanding debate on cortical columns as fundamental units of computation (Horton and Adams, 2005). The contrasting columnar structure we observed in V2 and V4, when viewed alongside the differences between these areas in terms of their size, inputs and physiological properties, sheds light on when columns might arise. V2 is the second largest visual area in the macaque brain, ∼80% the size of V1 (Vanni et al., 2020). V4 is much more compressed, about half the size of V2, yet it is considered to be the most interconnected visual cortical area in the macaque brain (Felleman and Van Essen, 1991). Functionally, V2 exhibits tuning for spatially homogeneous texture patterns, which may be computed by integrating basic features extracted from V1 (Freeman et al., 2013) and pooling neurons with similar feature preferences over spatial locations. In contrast, V4 processes more complex combinations of visual features, such as shape, texture, and color (Bushnell and Pasupathy, 2012; Okazawa et al., 2017; Kim et al., 2019) to build object-based representations (Pasupathy et al., 2020). The higher dimensionality of these representation, resulting from a convergence of diverse inputs onto V4, may require a more distributed coding strategy, leading to weaker spatial clustering (Purves et al., 1992). Thus, orientation-based columns in V1 are largely maintained in similarly sized V2. But a massive representational compression from V2 to V4, due to V4’s smaller size, combined with high input convergence onto V4 may prove to be unconducive for column formation.

### Properties of V2 encoding: implications for future studies

Prior studies have shown that V2 neurons encode combinations of orientations, such as angles and junctions (Ito and Komatsu, 2004; Anzai et al., 2007), possibly through the integration of inputs from V1 neurons with different orientation preferences. Consistent with this hypothesis, we found that V2 responses to shape stimuli could be largely modeled in terms of image characteristics as weighted combinations of orientation filter responses (Figure 4). This is quite unlike V4, where responses to shape stimuli cannot be explained in terms of a spectral receptive field model (Oleskiw et al., 2014). Instead, V4 responses reflect the characteristics of a shape boundary in object-centered coordinates. While future studies should confirm these findings with an expanded set of stimuli, our results indicate that a critical transformation may occur in the ventral visual pathway, where locally defined orientation combination in V2 are converted to globally defined, object-centered representations in V4 (Pasupathy and Connor, 2001).

With regards to motion direction selectivity, neurons in both V2 and V4 appear to encode information about drifting and translational motion. However, while V4 shows distinct tuning between these two motion types (Bigelow et al., 2023), V2 exhibits a moderate correlation between them. This raises the possibility that tuning for both motion types may arise from surface-based encoding in V2 (Adelson, 2001), whereas in V4, translational motion tuning may relate to object-based encoding (Pasupathy et al., 2020). Future studies could explore these V2/V4 differences in motion direction encoding and the underlying circuitry further. We also found that motion direction -selective neurons in V2 were not evenly distributed across recording sites but were instead concentrated at specific locations (Figure 5). This may relate to prior work linking V2 thick stripes to motion processing (Sincich and Horton, 2005; Lu et al., 2010).

Our analysis of temporal dynamics revealed that V2 neurons with sustained responses display stronger selectivity for both shape and texture features and are mainly distributed in superficial cortical layers. This laminar-specific pattern resonates with prior work suggesting that preference for naturalistic texture arises de novo in V2 (Ziemba et al., 2019). Future studies combining high-resolution anatomical techniques with neurophysiological recordings will be needed to study this process more precisely.

One limitation in directly comparing our current V2 results with previous V4 findings is the difference in experimental conditions. Specifically, the V4 data were collected from awake animals with a median of 5 repetitions per condition, while the current V2 data were obtained under anesthesia with 10 repetitions per condition. We consider it unlikely that anesthesia fundamentally alters the spatial clustering of similarly tuned neurons, but it might influence the magnitude, synchronization, and temporal dynamics of neuronal responses (Lee et al., 2021). Future studies using matched stimuli, recording methods, and animal states are needed to enable more rigorous comparisons across visual areas.

## Conclusion

Studies over decades have investigated the functional organization and the representational bases of various stages along the visual hierarchy, typically focusing on one area at a time with customized stimuli. Here, by comparing two successive visual areas with identical stimuli and recording methods, our results identify distinct properties of areas V2 and V4, reveal general insights into why columns might arise in the visual cortex, and offer constrains to improve future models of the ventral stream.

## Author contributions

T.K., T.N., and A.P. contributed to conception and design of the experiments. T.K., R.K., G.H., T.N., C.D., and W.B. conducted the anesthetized electrophysiology recordings. T.K., R.K., and G.H analyzed the data. T.K., R.K., G.H., and A.P. wrote the manuscript. W.B., and A.P. supervised the project and acquired funding. All authors approved the final version.

## Conflict of Interest

The authors declare no competing financial interests.

## Acknowledgements

The authors are grateful to lab members for providing helpful discussions and comments on the manuscript, and the WaNPRC veterinary staff and instrumental services for their support. This work was supported by NEI Grant R01 EY018839, EY029601 to A.P.; NINDS Grant U01 NS131810 to A.P. & W.B.; NEI Center Core Grant for Vision Research P30 EY01730 to the UW; NIH/ORIP Grant P51 OD010425 to the WaNPRC.

## Notes

### Competing Interest Statement

The authors have declared no competing interest.

