## Supplementary material for "Functional clusters for shape, texture, and motion encoding in macaque V2": Figure S1; Figure S2, Figure S3; Figure S4

### Supplemental Materials

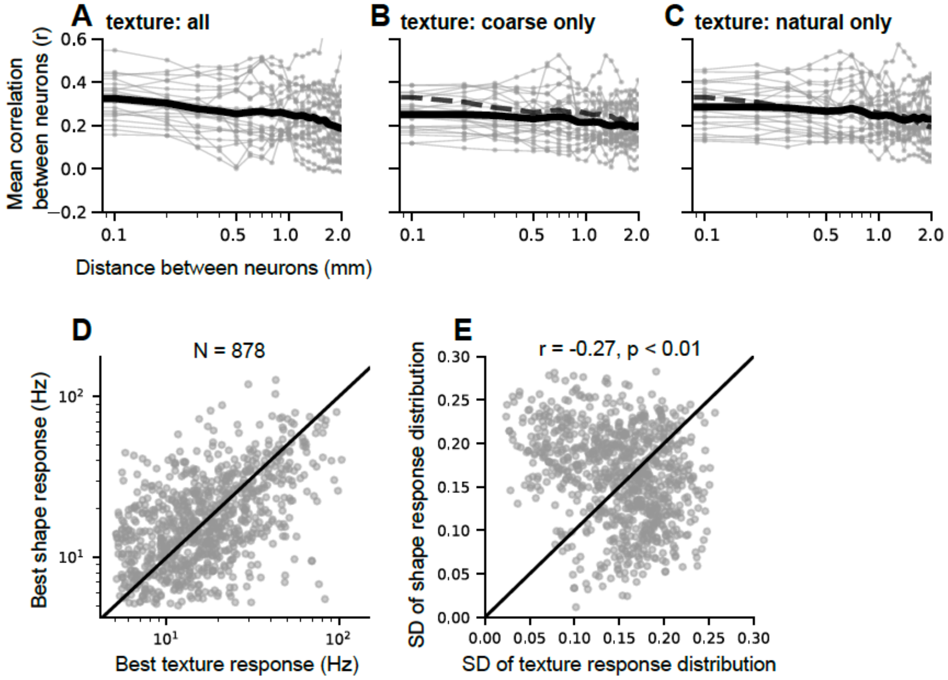


**Figure S1. Control analysis to validate larger neuronal clusters for textures. A.** Reproduction of the first panel in Figure 3B. Thin gray lines correspond to individual recording sites, with each data point representing the mean value of tuning similarity from 300 µm bins positioned every 100 µm. The black line indicates the average value across recording sites. **B.** The same analysis was conducted using coarse textures (excluding fine textures; see the main text for details). Dashed line shows data with all stimuli (as in **A**) for comparison. Excluding fine textures reduced response similarity between nearby neurons, but a substantially high similarity ($r > 0.2$) remained even at distances of ~1 mm. **C.** The same analysis was conducted using only natural textures (excluding noise textures). **D.** To test whether there is a notable difference in driven firing rates between texture and shape stimulus sets, the best shape and best texture responses from individual neurons were compared. Because these stimulus sets were tested in separate sessions (not simultaneously), to minimize potential effects of neuronal loss or emergence, only neurons exhibiting best responses greater than 5 spikes/s for both stimulus sets were included for this comparison (878 out of 1138 neurons). **E.** To compare the relative spread of responses for shapes and textures in individual neurons, responses were first normalized (1 representing the maximum response across both shape and texture stimuli) to control for firing rate differences, and then the standard deviation of each response distribution was calculated. Some neurons (dots below the diagonal line) exhibited greater response spread for textures, while others showed the opposite pattern. Overall, there was no strong bias toward either stimulus category.

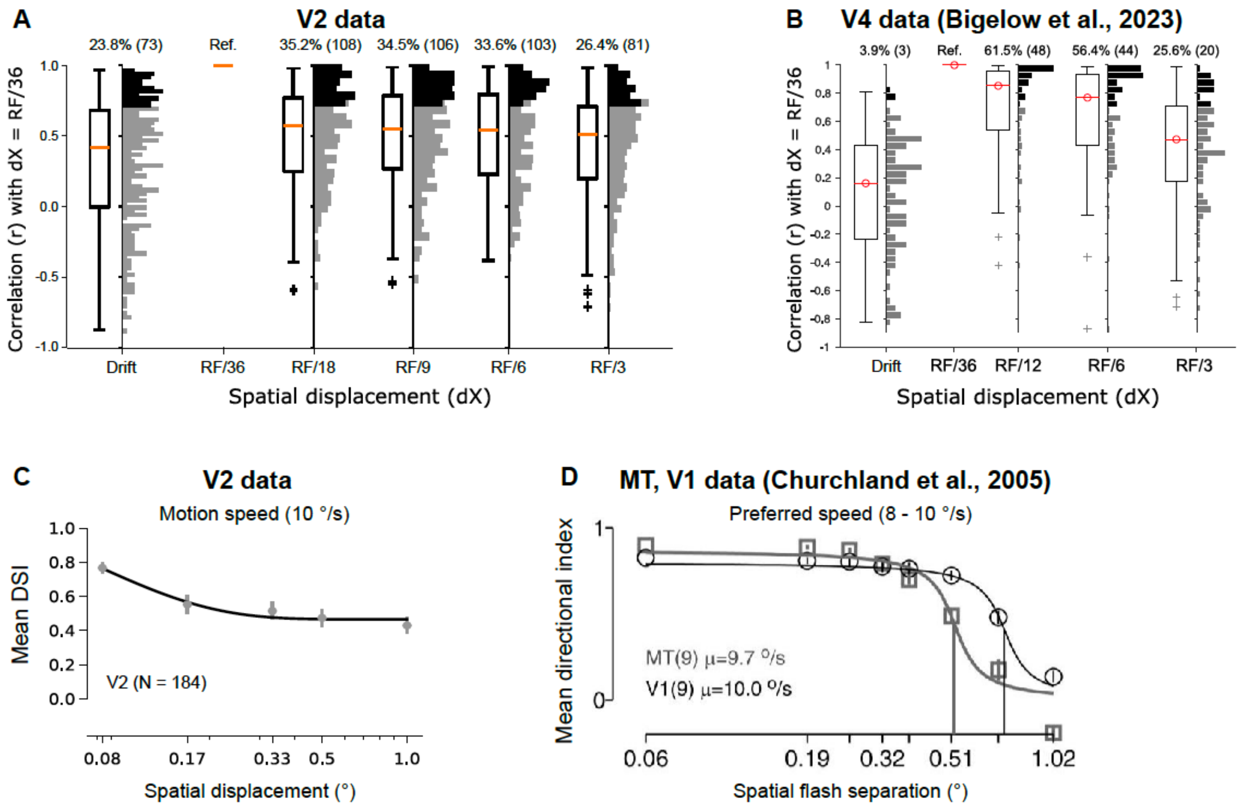


**Figure S2. Comparison of translational motion direction selectivity across visual areas. A.** Similarity between direction tuning curves from drifting and translational motion were assessed by computing the correlation between tuning curves for translational motion at dX = RF/36 and those for drifting motion and translational motion at other dX values. Top and bottom edges of each box display the interquartile range, and the red central mark is the median. The histogram alongside shows the entire distribution of values corresponding to each box plot. Filled bars indicate neurons with statistically significant correlation. Across the population, tuning curves showed substantial correlation (median $r > 0.4$) between tuning curves for short (RF/36) and long (RF/3) spatial displacement conditions, and drifting and translational motion. **B.** Figure reproduced from (Bigelow et al., 2023), with the identical analysis applied to V4 as shown in **A.** Unlike V2, tuning curves between drifting and translational motion (the first box plot) were very weakly correlated in V4. **C-D.** Direction selectivity for translational motion as a function of spatial displacement. V2 direction selectivity remained moderate even at 1° displacement, whereas direction selectivity is not evident at this level in area V1 and MT. **D.** adapted from (Churchland et al., 2005), showing data from V1 and MT.

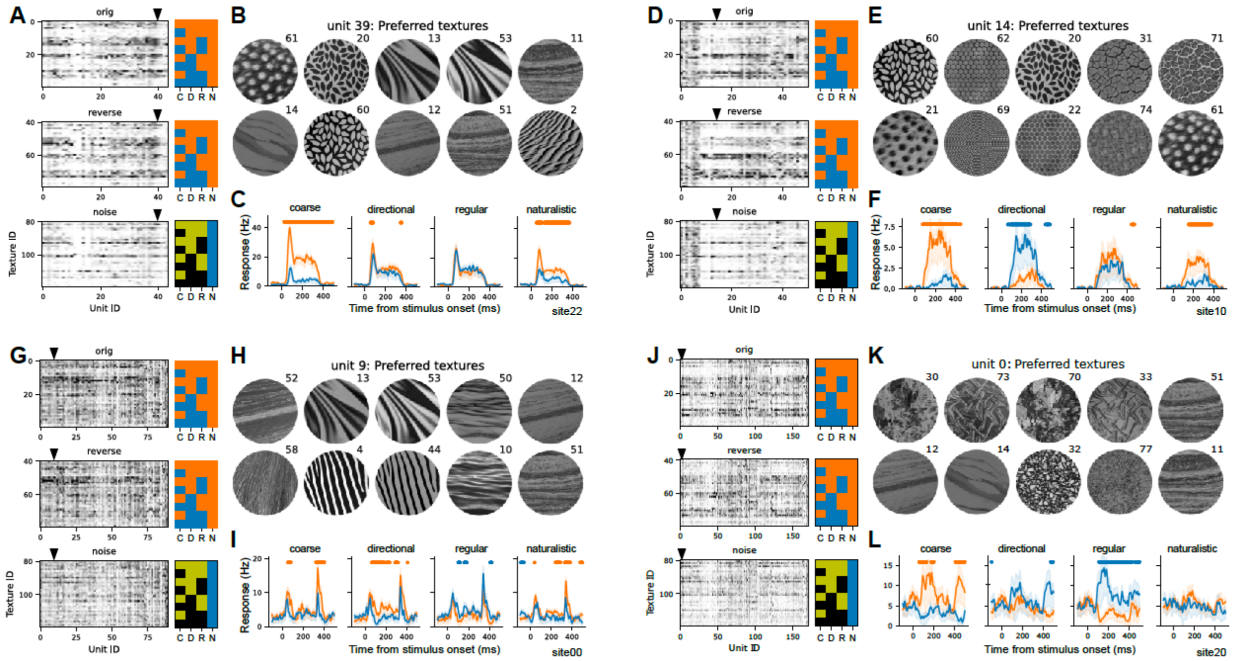


**Figure S3. Neuronal clusters with different texture selectivity.** **A.** Reproduction of Figure 6A. **B.** For a specific neuron (unit 39) from this site, the 10 most preferred textures based on the spike count during the 0–400 ms window after stimulus onset are shown. These textures include a mix of directional and non-directional (#20, 60, 61) patterns, as well as regular (#2, 20, 60, 61) and irregular structures, but all belonging to the coarse category. **C.** The mean PSTHs for two opposing levels of texture dimensions are displayed in orange and blue, respectively. Shaded areas indicate ± one standard error of the mean. Dots above the lines indicate time bins in which a statistically significant difference between the mean responses was observed. This example neuron demonstrates a strong preference for coarse and naturalistic features, with minimal sensitivity to directionality or regularity. **D-F.** The same representation as in **A-C** for an example neuron at site 10 (Figure 6C-D). The 10 most preferred textures include both regular and irregular (#31, 71, 74) patterns, but all are non-directional and nine (all except texture #69) are coarse (**E**). This multi-dimensional texture selectivity is also evident in the temporal response dynamics (**F**). **G-I.** Results from an example neuron at site 00 (Figure 6E-F). In contrast to the unit in **D-F**, the top ten preferred textures are all directional. These include both regular (#4, 44) and irregular patterns. **J-L.** Results from an example neuron at site 20 (Figure 6G-H). The 10 most preferred textures are all irregular, and nine (all except #77) are coarse. These preferences span both directional (#11, 12, 14, 51) and non-directional patterns. Interestingly, the neuron's responses were enhanced by coarse, irregular textures and suppressed by fine, regular textures.

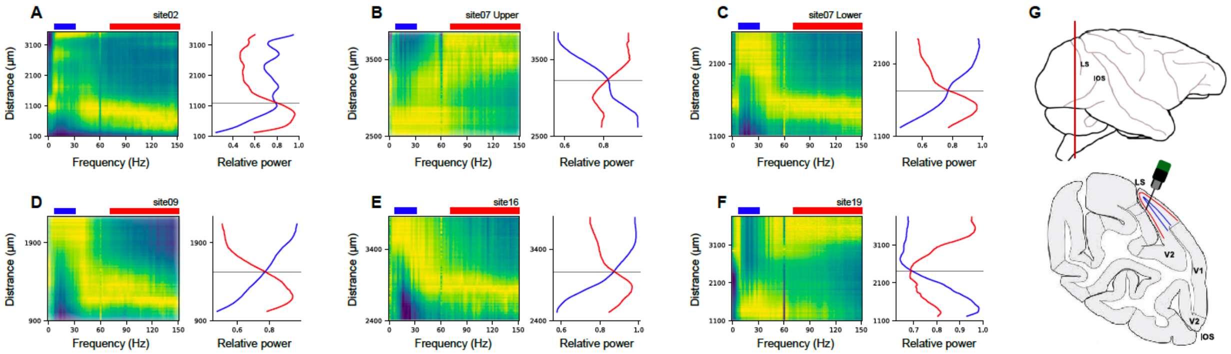


**Figure S4. Example spectrolaminar patterns.** To estimate the positions of probe channels across cortical layers, we calculated and compared the relative LFP power in the alpha-beta (blue; 10-30 Hz) and gamma (red; 75-150 Hz) frequency bands (Mendoza-Halliday et al., 2024) (see Methods). The crossover between these two power bands (indicated by the gray horizontal line) was used to estimate the location of layer 4. Higher gamma power relative to alpha-beta power was considered indicative of superficial layers, while the opposite was considered indicative of deeper layers. Panels **B** & **F** show spectrolaminar patterns when the probe crossed layers in superficial-to-deep direction (with the deep layers near the probe tip). Due to the anatomical characteristics of area V2 (**G**), when the probe is positioned deep within the posterior bank of the lunate sulcus, inverted spectrolaminar patterns are often observed. In these cases, the transition runs from deep to superficial layers, with superficial layers represented by channels closer to the probe tip (**A, C, D, E**).
